# MyeloDB: A multi-omics resource for Multiple Myeloma

**DOI:** 10.1101/2023.05.18.541396

**Authors:** Ambuj Kumar, Keerthana Vinod Kumar, Kavita Kundal, Avik Sengupta, Kunjulakshmi R, Rahul Kumar

## Abstract

Multiple myeloma (MM) is a common type of blood cancer affecting plasma cells originating from the lymphoid B-cell lineage. It accounts for about 10% of all haematological malignancies and can cause significant end-organ damage. The emergence of genomic technologies such as next-generation sequencing and gene expression analysis has opened new possibilities for early detection of multiple myeloma and identification of personalized treatment options. However, there remain significant challenges to overcome in MM research, including integrating multi-omics data, achieving a comprehensive understanding of the disease, and developing targeted therapies and biomarkers. The extensive data generated by these technologies presents another challenge for data analysis and interpretation. To bridge this gap, we have developed a multi-omics open-access database called MyeloDB. It includes gene expression profiling, high throughput CRISPR-Cas9 screens, drug sensitivity resources profile, and biomarkers. MyeloDB contains 47 expression profiles, 3 methylation profiles comprising a total of 5630 patient samples and 15 biomarkers which were reported in previous studies. In addition to this, MyeloDB can provide significant insight of gene mutations in MM on drug sensitivity. Furthermore, users can download the datasets and conduct their own analyses. Utilizing this database, we have identified five novel genes i.e., *CBFB, MANF, MBNL1, SEPHS2* and *UFM1* as potential drug targets for MM. We hope MyeloDB will serve as a comprehensive platform for researchers and foster novel discoveries in MM. MyeloDB is freely accessible at: (https://project.iith.ac.in/cgntlab/myelodb/)

## 1. Introduction

Multiple myeloma (MM) is one of the most prevalent blood cancer and accounts for around 10% of all blood malignancies; it is diagnosed by the presence of abnormal clonal plasma cells in the bone marrow (1). Studies have shown that patients with multiple myeloma, come with anaemia (73%), osteolytic bone disease (79%), and acute kidney injury (19%) (2). MM is a type of cancer affecting plasma cells originating from the lymphoid B-cell lineage (3). Asymptomatic premalignant plasma cell proliferation gives rise to MM. These plasma cells are called “Monoclonal Gammopathy of Undetermined Significance” (MGUS) (4). In patients where MM is asymptomatic, it is called “Smoldering multiple myeloma (SMM)”, which later transforms into malignant MM causing end-organ damage (4).

GLOBOCAN 2020 data shows a worldwide estimate of 1,76,404 cases of MM with an estimated mortality of 1,17,077 cases (5). MM has a lifetime risk of 1 in 132 (0.76%) (6). There has been an increased understanding of this disease by knowing precursor conditions, which has helped in developing new strategies to tackle MM, including next-generation agents (antimyeloma drugs) and immunotherapies (7).

According to the data, 37% of patients diagnosed with the MM were under the age of 65, and the median age at diagnosis was between 66 and 70 years. In contrast, cases of the MM occurring in patients under 30 years old are extremely rare, with reported frequencies ranging from 0.02% to 0.3%, and the MM appears to be more prevalent in men in this age group (8–11). Reports show that over 50% of patients with MGUS have had MM for over ten years, but due to the asymptomatic nature of MGUS, the diagnosis of MM at this stage is difficult. MGUS progresses at a rate of 1% per year, and 3% of the population aging above 50 has MGUS (12–16). An advanced stage of MGUS, which is also asymptomatic, called SMM, can progress to MM at a rate of 10% per year (first five years following diagnosis), 3% per year over the next five years, and 1.5% per year after that (17,18). Data suggests that the following cytogenesis of the disease influences this progression rate, such as a patient with translocation t(4;14), del(17p), and gain(1q) having more risk of progressing from SMM to MM (19).

MM research is currently emphasizing how the disease develops from precursor conditions, the utilization of minimal residual disease as a prognostic tool, and the rapid advancement of new treatment options (7). These treatments include next-generation versions of existing antimyeloma drugs and novel therapies with different mechanisms of action, particularly new immunotherapies (7).

Experts are exploring different ways to develop new biomarkers to improve diagnosis, prognosis, and treatment. Identifying effective, sensitive, and specific biomarkers is crucial for detecting the disease at its preliminary stages. Several ongoing studies are underway to investigate novel biomarkers for the diagnosis and prognosis of myeloma (20). Due to advancements in our understanding of the biology of multiple myeloma, novel diagnostic and prognostic methods have been developed (20). Understanding the disease’s pathogenesis and the molecular concept behind these novel biomarkers is crucial to ensure their validation and correct usage (20). With the progress in genomics, researchers produce enormous amounts of data to examine the transcriptomic level expression of both cancerous and normal cells and gain insight into the pathology of the disease. Previously, researchers have conducted numerous studies using high-throughput techniques to identify biomarkers specific to cancer. Most of the data generated by these high-throughput techniques have been stored in various databases over time, such as the Genomic Data Commons (GDC) data portal (21), Genomics of Drug Sensitivity in Cancer (GDSC) (22), the International Cancer Genome Consortium (ICGC) Data Portal (23), DepMap (Dependency map) (24), The Cancer Genome Atlas (TCGA) (25), and the Gene Expression Omnibus (GEO) (26). One of the major drawbacks of these databases is variation in the file format of expression datasets with most of the data present in a raw format. Another challenge is the lack of a centralized repository of patient data, which can help in the identification of prognostic factors and guide personalized treatment decisions. This fact makes it difficult and time-consuming for the researcher to analyse and extract biologically relevant information.

To the best of our knowledge, there is no database of MM that can overcome this shortcoming of existing database. The scattered data on MM hinders the development of a comprehensive understanding of the disease and impedes advancements in the field. To fill the information gap, we present a MM database called “MyeloDB”, a multi-omics database of gene expression profiling, high throughput CRISPR-Cas9 knockout screens, drug sensitivity resources profile, and biomarkers. MyeloDB aims to provide scattered information on a single platform in a homogenized format. MyeloDB offers data of 5630 expression profile samples and 231 methylation profile samples, and 17387 CRISPR and 18010 Achilles genes found in multiple myeloma. MyeloDB also has drug sensitivity profiles of 17 MM cell lines. Users can access this informative data through user-friendly interfaces and unravel the mysteries of MM. MyeloDB is a powerful and efficient database with credible content, which can provide improved and precise information for gene targets and biomarker research in multiple myeloma (MM). Users can access MyeloDB at https://project.iith.ac.in/cgntlab/myelodb.

## 2. Materials and Methods

### 2.1 Data Collection

#### 2.1.1 Gene expression profile dataset

We used the following keywords and criteria to obtain gene expression profiles from Gene Expression Omnibus (GEO): “multiple myeloma”. GEO datasets were filtered using the following criteria: a) organism: *“Homo sapiens”*, b) study ‘type’: “expression profiling by array”, and “expression profiling by high throughput sequencing”. Furthermore, we exclude studies based on cell lines and datasets with less than 6 samples. We included GEO datasets from sub-series and excluded their corresponding super-series to eliminate data redundancy. MyeloDB includes datasets that are limited to studies published before January 2023. Initially, around 733 expression profile datasets were present for MM, including profiling by arrays and high throughput sequencing. Data was categorized and processed based on their profiling techniques.

We searched the datasets for information like “Gene Symbol” or “Entrez ID” and “Probe ID” after obtaining normalized expression values from different profiling techniques. We downloaded the relevant data for further processing, and we excluded datasets that were unavailable for download or had truncated data, such as data that could not be read or data that was missing. As a result, we ended up with 47 gene expression datasets comprising 5630 samples.

#### 2.1.2 Gene methylation profile dataset

To obtain methylation profile datasets following criteria were used to search for profiles: a) organism: *“Homo sapiens”*, b) study ‘type’: “methylation profiling by array,” and “methylation profiling by high throughput sequencing”. Only 17 datasets were present in GEO on or before January 2023. After filtering datasets with “study type” as *Homo sapiens* and removing cell lines data, 10 datasets remained. Furthermore, we downloaded data using the *GEOquery* package in R for raw and supplementary files. Datasets with information like “Gene Symbol” or “Entrez ID” and “Probe ID” were downloaded for further processing, and we did not consider datasets that were unavailable for download or had truncated data, such as missing values or unreadable information. At last, 3 methylation datasets remained with a total of 231 samples.

#### 2.1.3 Knockout CRISPR-Cas9 screens

We used the DepMap portal (https://depmap.org/portal/) (24) to download CRISPR-Cas9 based knockout screen and project achilles data for MM. DepMap public 22Q2 was accessed and files ‘CRISPR_gene_effect.csv’, ‘Achilles_gene_effect.csv’, and ‘Sample_info.csv’ were downloaded. We filtered the Sample_info file for primary disease as ‘myeloma’ and subtype as ‘Multiple Myeloma’, which resulted in 34 MM cell line datasets. These 34 cell lines were mapped with CRISPR and Achilles datasets using their DepMap ID. We found 20 cell lines for MM, which are included in our MyeloDB database.

#### 2.1.4 Drug sensitivity data

Genomics of Drug Sensitivity in Cancer (GDSC) (https://www.cancerrxgene.org/) (22) database stores 969 cell lines, 297 drug compounds, and 243466 IC50s in GDCS2. We downloaded the GDSC2 data from the GDSC database. While downloading drug dataset, “screening set” was selected to “GDSC2”; “Target pathways” as “all” and “Tissue” as “multiple myeloma”. We filtered 17 cell lines that were classified as MM according to TCGA and had a tissue subtype specified as myeloma. 279 drug compounds with IC50 data against 17 MM cell lines were available from GDSC. This drug sensitivity data of 17 MM cell lines along with mutation data is used for the mutational analysis.

#### 2.1.5 Mutation data

Mutation data was downloaded from COSMIC (Catalogue of Somatic Mutations in Cancer) database (https://cancer.sanger.ac.uk/cell_lines) (27). We used Cell Lines Project v97 (released on 29-NOV-22) to download the complete mutation data (CMD) file. Out of the 55009 unique gene mutations in CMD file, “Plasma cell myeloma” was filtered to get MM data. We further filtered this data to include only the 17 MM cell lines obtained from GDSC. As a result, we found 32381 gene mutations for 17 MM cell lines in CMD file. We further removed mutations described as “unknown”, “substitution-coding silent”, or “nonstop”. This resulted in a final dataset of 12,025 unique gene mutations in 17 MM cell lines. This dataset was merged with the drug sensitivity dataset and used for the mutational analysis in MyeloDB.

### 2.2 Data processing

We downloaded the raw files for expression and methylation profiling data using the *GEOquery* package in R. Each raw file passed through a detailed curation process. Datasets had different file formats such as CEL, TSV, CSV, xlsx, and txt, so it was necessary to read and make them in a single file format that researchers could access easily. We developed individual pipelines for Affymetrix, Illumina and Agilent datasets. Figure 1 shows the workflow of MyeloDB.

a. Affymetrix: We individually read the Affymetrix expression profiles and normalized them using RMA (Robust Multichip Average) normalization method in Bioconductor package *affy* (28,29) or *oligo* (30). These data frames then were merged with clinical data obtained from corresponding matrix file of each dataset. Clinical data included the sample source, patient age/sex, sample type, etc. We generated a final matrix file for all datasets in CSV format, which are available for download on MyeloDB. For some expression datasets, raw data was not available, so we separately read them and applied the background correction and quantile normalization. Then, we generated a matrix file and included it in MyeloDB.
b. Illumina: We used the *limma* package (31,32) to process Illumina data. For expression microarray data, we downloaded non-normalized background-corrected data from GEO and converted it to quantile-normalized values. The Illumina high-throughput sequencing data containing raw count, RPKM (Reads per kilobase of transcript per Million reads mapped), or FPKM (Fragments Per kilobase of transcript per Million reads mapped values) were obtained and subsequently transformed into transcripts per million (TPM) values to facilitate downstream analysis.
c. Agilent: For Agilent expression profiling data, we downloaded raw files using the *GEOquery* package and then background-corrected and quantile-normalized the values. These expression values were merged with the clinical data which we downloaded using getGEO() in R. Finally, a matrix file in CSV format is generated which can be downloaded through our database.
d. Methylation data: In case of Illumina methylation array data, we retrieved beta values from the raw/supplementary files or series matrix files using function getGEO() in R. These expression/methylation values were merged with clinical characteristics data from series matrix files. We generated a final matrix file for all datasets in CSV format and incorporated in MyeloDB.

**Figure 1.**
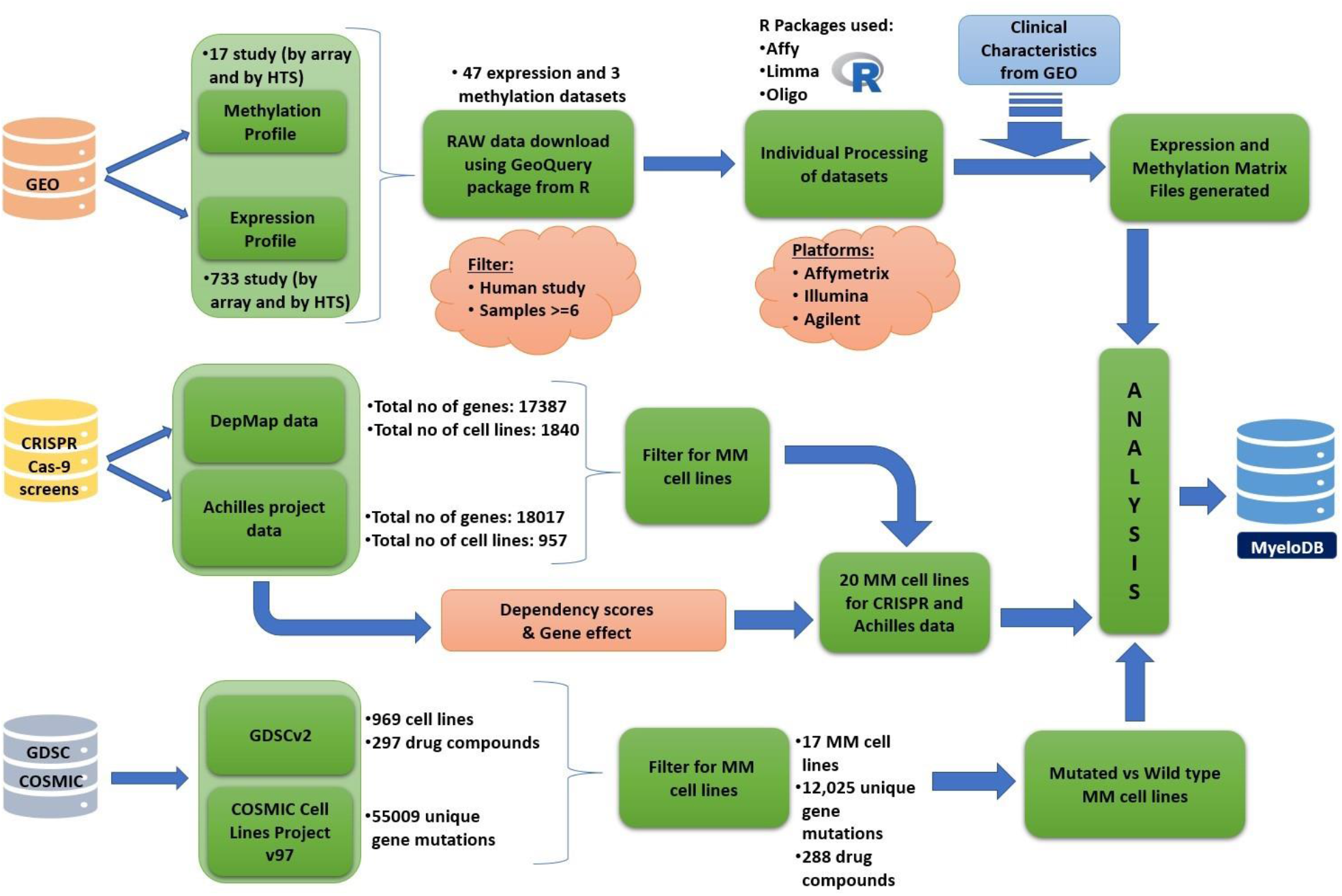
Workflow of MyeloDB.

In our generated matrix file, if only the Entrez ID was present, we used R to retrieve the corresponding gene symbols and added them to the appropriate matrix files. We generated a final matrix containing clinical information and made it available for use in MyeloDB.

### 2.3 Database and website implementation

MyeloDB is built on an Ubuntu-based machine’s Apache HTTP Server (v2.4.54). The front end was created using HTML5, CSS3, PHP5, and JavaScript and it is designed for mobile, tablet, and desktop compatibility. The back end utilizes MySQL/10.4.25-MariaDB to manage data.

## 3. Results & Discussion

### 3.1 Database statistics

MyeloDB is a resource having 47 expression profiles of 5630 samples, 3 methylation profiles of 231 samples, drug sensitivity and mutation data for 17 MM cell lines, CRISPR-Cas9 knockout screens data from DepMap (17387 genes) and Achilies project (18017 genes) dependency scores for 20 MM cell lines.

### 3.2 Website interface and contents of MyeloDB

MyeloDB is a freely accessible at (https://project.iith.ac.in/cgntlab/myelodb/) with a user-friendly interface designed to operate optimally with Google Chrome and Firefox browser. The MyeloDB website offers users an intuitive browsing and searching experience, with seven main functional pages: ‘Home’, ‘Datasets’, ‘Analysis’, ‘Biomarker’, ‘Download’, ‘Help’ and ‘Team’. The website’s structure is depicted in Figure 2.

I. The Home page provides users with MyeloDB statistics, contents, and database workflow.
II. The ‘Datasets’ tab aims to provide users access to various publicly available data repositories. We divided the “Dataset” tab into 3 sections according to data source: “GEO data”, “CRISPR-cas9 screens”, and “GDSC data”.
  A. GEO data: it comes with the option of “browse on” which lets the user search query according to different fields such as profiling technique, GSE ID, PubMed ID, Sample source, etc. It also has a tab that lets users browse datasets using the “profiling technique”.
  B. CRISPR-cas9 screens data: This section allows the user to search gene dependency information of 17387 genes (DepMap) and 18017 genes (Achilles project) from in 20 MM cell lines from DepMap portal. User can also get this information in graphical format by clicking plot button.
  C. GDSC data: it has a “Browse by Drug” option which allow users to search a drug (279 drug data is stored in MyeloDB) and it returns with IC50 values of the respective drug in MM cell lines.
III. The ‘Analysis’ section provides users an access to the analyses we have conducted using the MyeloDB data. The analysis option has two sections, “Drug sensitivity “ and “profiling analysis”.
  A. Drug sensitivity: This section allows users to select one gene and one drug to identify the effect of gene mutation on drug sensitivity.
  B. Profiling analysis: This section allows users to search and visualize a gene from expression and methylation profiles from GEO data.
IV. The “Biomarker” option allows user to get data of MM biomarkers reported in the literature from PubMed. It provides information on biomarkers’ type, source, and regulation with the significance and methods used to identify them.
V. The “Download” option allows users to download the entire datasets available in MyeloDB.
VI. The “Help” section assists users in browsing the website and provides a glossary to familiarize them with usage of MyeloDB.
VII. The “Team” section acknowledges the contributions of researchers behind this database.

**Figure 2.**
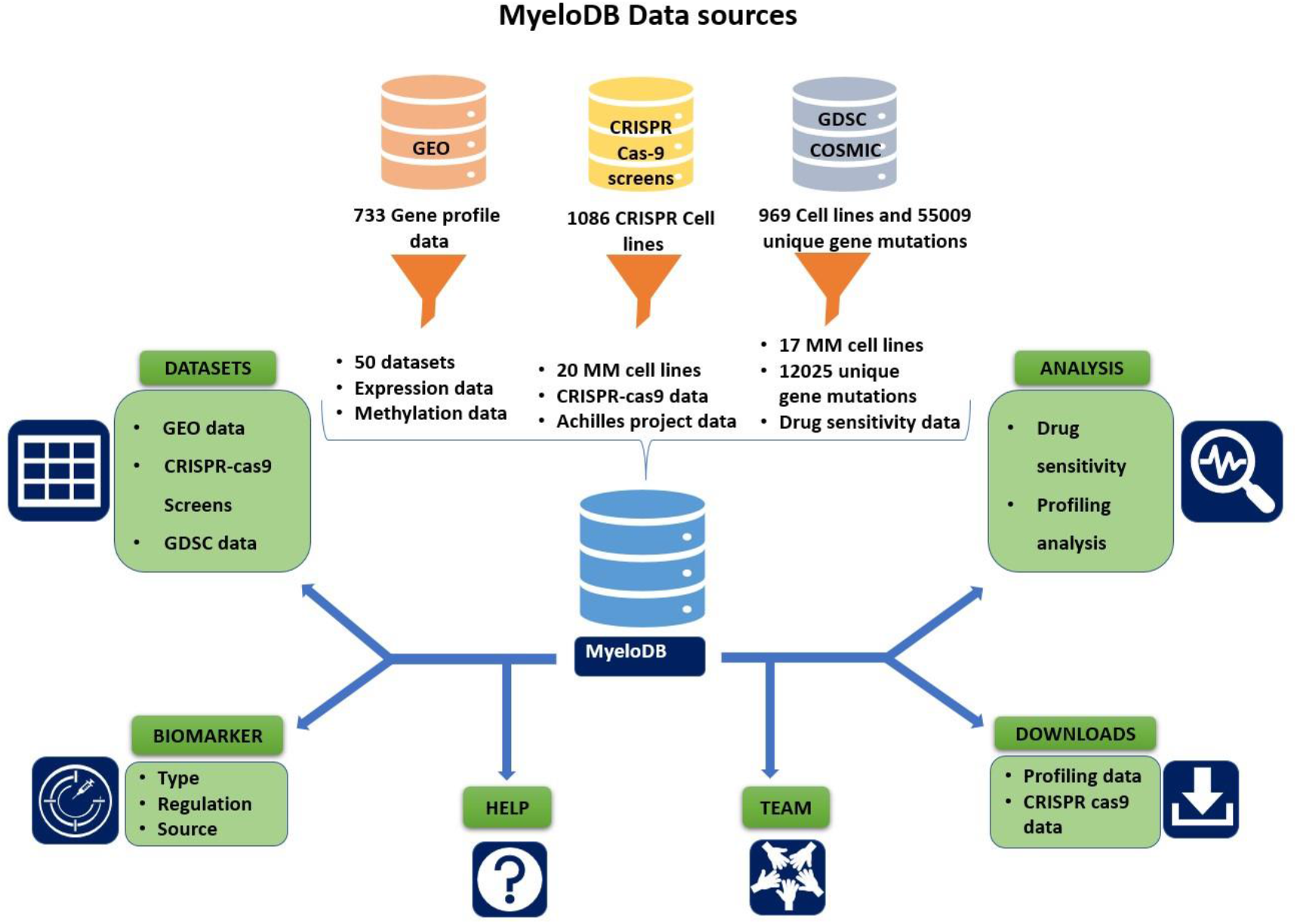
MyeloDB Website structure.

### 3.3 Analysis

I. **Mutational analysis:** The mutational analysis section of drug sensitivity allows users to search a gene and a drug pair, which will provide a table with information on mutated and wild-type cell lines of MM for searched gene and IC50 data for searched drug against these cell lines. In addition to this user can also visualize the result in box plot comparing the IC50 for mutated and wild-type cell lines with p values. p < 0.05 is considered as significant. Figure 3(A) and 3(B) shows two most significant gene-drug combination where mutation in *CCDC80* and *HMCN1* has sensitized MM cell lines against the drug Entinostat and I-BET-762 respectively. Figure 3(C) and 3(D) shows two most significant gene-drug combination where mutation in gene *LAMA2 and TP53* has made cell line resistant against the drug EHT-1864 and CDK9_5576 respectively.
II. **Profiling analysis:** This section allows users to analyse gene expression and methylation patterns in diverse types of profiling techniques i.e., Affymetrix, Illumina, and Agilent. We provide users with a search option for a gene, which returns the mean expression or methylation values across all the techniques. In addition, users can visualize the plots to better understand the mean expression values of genes. User can download these plots for each technique and utilize them for their research.

**Figure 3.**
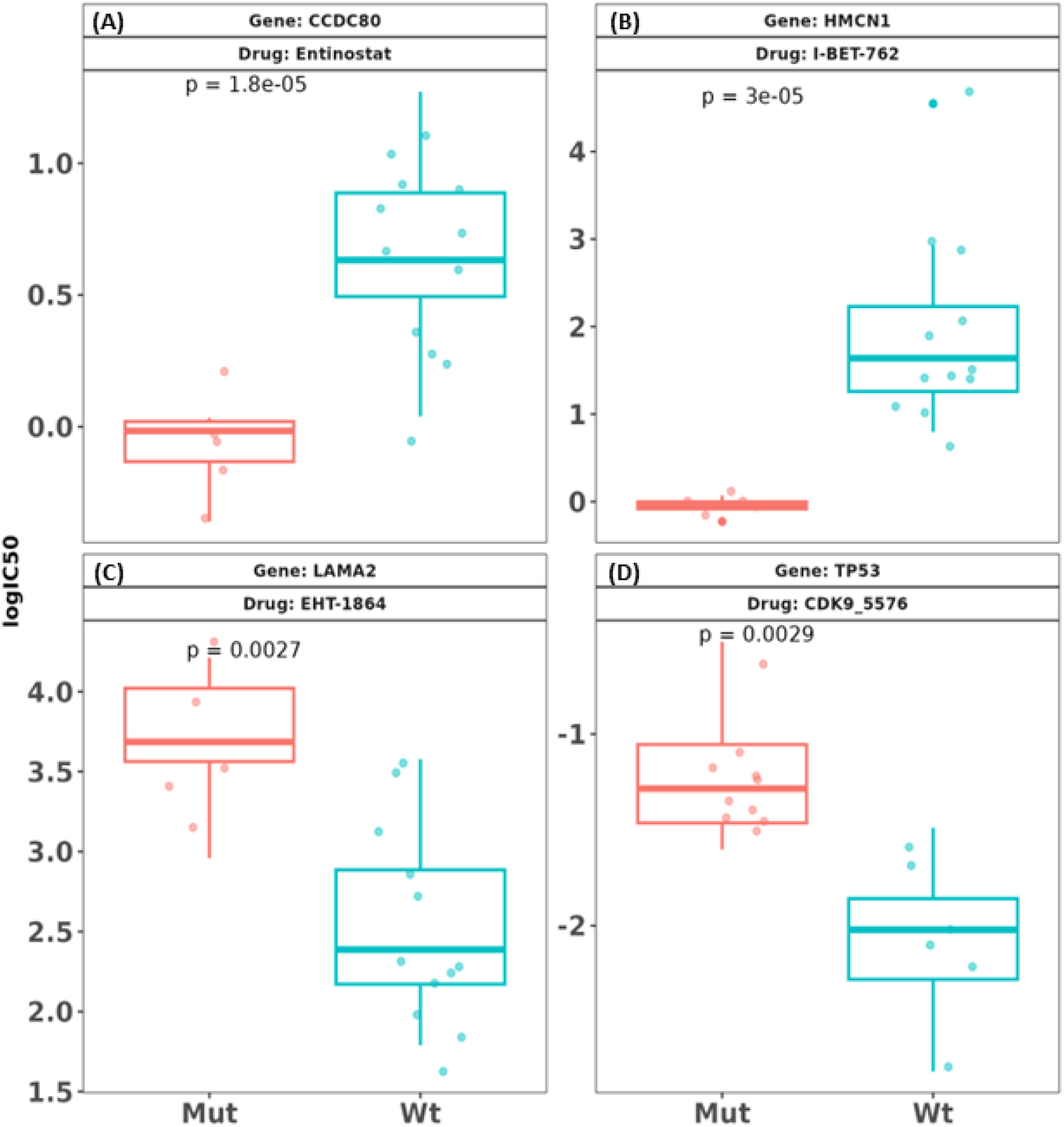
Mutation in genes (A) *CCDC80 and* (B) *HMCN1* sensitize MM cell lines for the drugs Entinostat and I-BET-762 respectively. Mutation in genes (C) *LAMA2* and (D) *TP53* decrease the sensitivity of drugs EHT-1864 and CDK9_5576 for MM cell lines respectively.

### 3.4 Utilizing MyeloDB for potential gene target identification

We used Affymetrix expression profile datasets and gene dependency data from MyeloDB. This integrative analysis guided us to identify novel genes as potential drug targets for MM. In this analysis we used Affymetrix datasets having 30 (RMA) and 7 (quantile normalized) datasets separately. To obtain the highly expressed genes, we computed the mean expression values of each gene across all tumor samples in each dataset and selected the top quantile genes (assuming these are over-expressed in MM). Next, we excluded housekeeping genes (33) and combined the remaining genes with CRISPR-Cas9 knockout screens based gene dependency scores. We then counted for the number of cell lines, where dependency scores were more than 0.7. We filtered the genes with counts greater than or equal to 10 (i.e., considering that 50 percent of MM cell lines depend on these genes for survival). We applied this pipeline to each of the datasets of Affymetrix and filtered out the common genes. As a result, we got 64 genes common in RMA normalized datasets and 61 genes common in quantile normalized datasets. Lastly, we eliminated the genes, which are common essential across different cancer types as provided in DepMap portal. Finally, we identified 4 genes from the RMA datasets i.e., *MANF, SEPHS2, CBFB* and *UFM1*, while we identified 7 genes from the quantile normalized datasets i.e., *MANF, CBFB, CCND2, MYH9, PIM2, MBNL1 and MEF2C*. Interestingly, of these 9 unique genes (*MANF* and *CBFB* are common in both) and 4 genes (*CCND2* (34), *MEF2C* (35), *PIM2* (36) and *MYH9* (37) have already been reported to have direct or indirect association with MM. The CCND2 gene has been identified as being overexpressed in multiple myeloma (MM). Furthermore, researchers have discovered and validated a super-enhancer that regulates the expression of *CCND2* in MM patients (34). Another essential gene for MM cell proliferation and survival is *MEF2C*, which has been reported to be overexpressed at the RNA level. It has been suggested that *MEF2C* could serve as a potential therapeutic target in MM (35). *PIM2*, known for its expression in a wide range of solid and hematological malignancies, has also been found to be overexpressed in MM patients (36). Differential expression analysis between MGUS and MM revealed *MYH9* as a significant gene and it is involved in shape change function, highlighting its role in MM (37).

## 4. Conclusion

MyeloDB is a powerful resource integrating various multi-omics studies across publicly available data portals. It can be a valuable resource for identification of novel and potential therapeutic targets for MM. Figure 4 shows the applications of MyeloDB. Microarray and RNA-Seq technologies have led to a significant surge in multi-omics data for MM over the years. But this data is scattered among different platforms with varying formats, which hinders its optimal usage and slows down the productivity and efficiency of MM research. The lack of uniform data matrices in an adequate format often leads to a biased interpretation of the biological significance of these data. To fill this gap, we developed MyeloDB, an integrative resource with expression and methylation profiles, CRISPR-Cas9 based knockout screens and drug sensitivity data and its association with gene mutation in MM. From our analysis using MyeloDB, we reported 9 genes as potential drug targets against MM. Of these 9 genes, 4 genes have been shown previously associated with MM and 5 genes are reported for the first time in our analysis. Our analysis warrants further investigation on these genes for their association with MM pathology. In addition to this, users can also download the datasets and perform their own analysis. We anticipate that MyeloDB will become an indispensable platform for researchers and facilitate the understanding of MM pathology.

**Figure 4.**
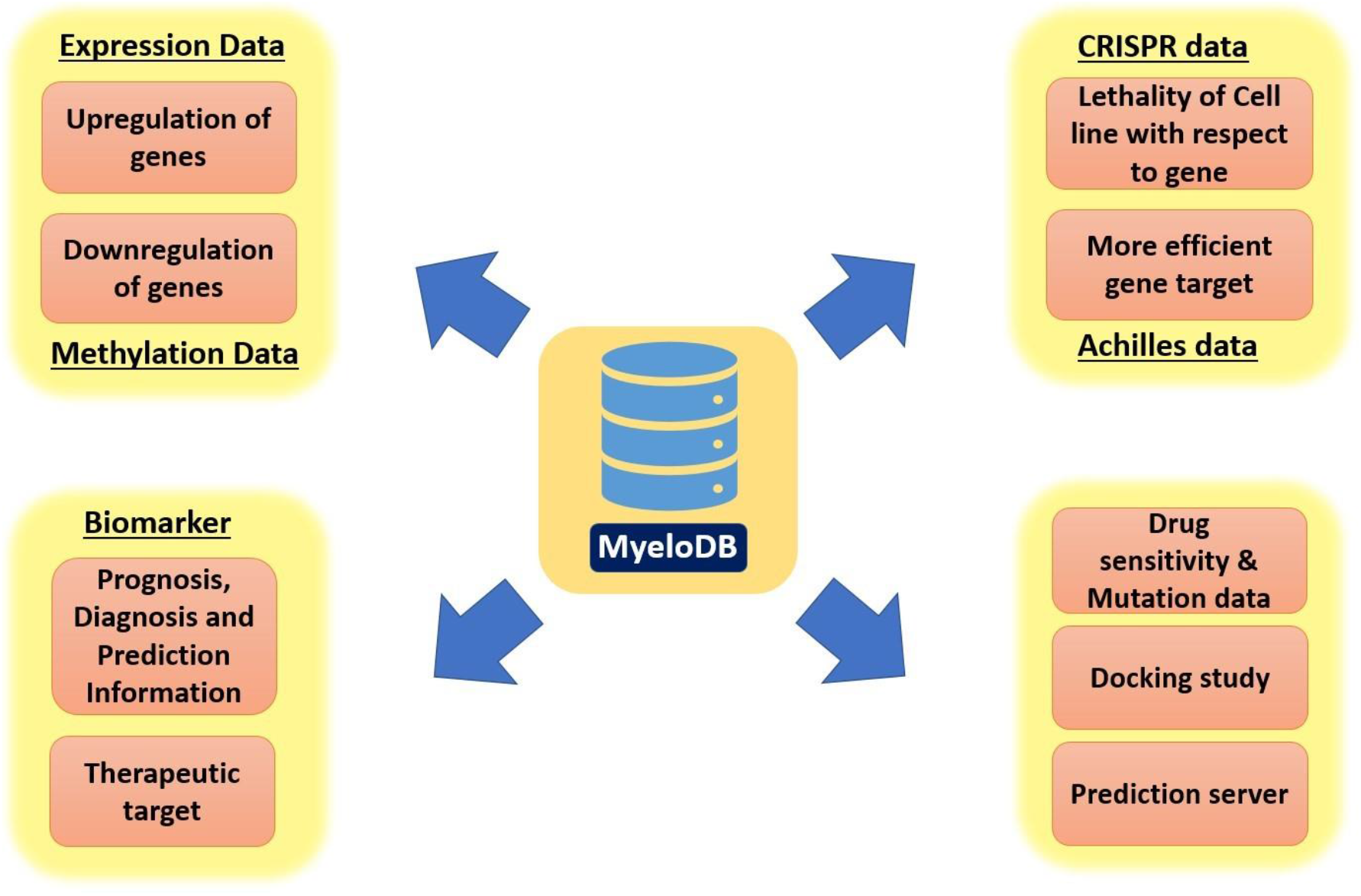
Applications of MyeloDB

## 5. Author Contribution

AK, KVK and RK conceive the idea and prepare the database framework. AK and KVK collected and analysed the datasets. AK developed the database and wrote first draft of the manuscript. KK, AS and KR helped in collecting the datasets and preparing the database outline. All authors read the manuscript and approved it for submission.

## 6. Conflict of Interest

Authors declare no conflict of interest.

## 7. Acknowledgements

AK, KVK and KK acknowledge the fellowship from Ministry of Education, Government of India. AS acknowledge the fellowship from Council of Scientific and Industrial Research (CSIR), Government of India. We are thankful to the Indian Institute of Technology Hyderabad for providing research infrastructure.

## Notes

### Competing Interest Statement

The authors have declared no competing interest.

